# Automation of tree-ring detection and measurements using deep learning

**DOI:** 10.1101/2022.01.10.475709

**Authors:** Miroslav Poláček, Alexis Arizpe, Patrick Hüther, Lisa Weidlich, Sonja Steindl, Kelly Swarts

**Affiliations:** Gregor Mendel Institute, Austrian Academy of Sciences, Dr. Bohr-Gasse 3, 1030, Vienna, Austria; Department of Forest Ecology, Faculty of Forestry and Wood Sciences, Czech University of Life Sciences, Kamýcká 129,165 00, Prague, Czech Republic; Faculty of Biology, Ludwig-Maximilians-University Munich, 82152 Martinsried, Germany; Max Perutz Labs, Department of Structural and Computational Biology, University of Vienna, Campus-Vienna-Biocenter 5, 1030, Vienna, Austria

**Keywords:** automation, computer vision, mask R-CNN, phenotyping, tree rings

## Abstract

1. We present an implementable neural network-based automated detection and measurement of tree-ring boundaries from coniferous species.
2. We trained our Mask R-CNN extensively on over 8,000 manually annotated rings. We assessed the performance of the trained model from our core processing pipeline on real world data.
3. The CNN performed well, recognizing over 99% of ring boundaries (precision) and a recall value of 95% when tested on real world data. Additionally, we have implemented automatic measurements based on minimum distance between rings. With minimal editing for missed ring detections, these measurements were a 99% match with human measurements of the same samples.
4. Our CNN is readily deployable through a Docker container and requires only basic command line skills. Application outputs include editable annotations which facilitate the efficient generation of ring-width measurements from tree-ring samples, an important source of environmental data.

## Introduction

Annual growth measurements estimated from tree increment cores are used in climate studies (Fritts, 1971), ecology (Primicia et al., 2015) and, increasingly, breeding (Housset et al., 2018). Tree-rings are traditionally hand-measured (Larsson, 2003), a time- and human-intensive process that inhibits the generation of very large datasets. Also, large datasets or meta-analyses may be biased when multiple people contribute to measurements. Analysis on digital images of tree-ring samples is becoming increasingly common, as are developments in both image acquisition systems (Griffin et al., 2021; Levanič, 2007) and programs for analysis (Crawford, 2021; Rademacher et al., 2021)). Additionally, some forms of dendrochronological analysis that depend on high resolution images such as blue intensity (Rydval et al., 2014) or quantitative wood anatomy (von Arx & Carrer, 2014) require digital images.

CNNs are widely used across many fields from facial recognition (Yang & Yin, 2017) to human health (Kido et al., 2018), finance (Gudelek et al., 2017), genomics (Washburn et al., 2019) and ecology (Fricker et al., 2019). Increased use of imaged increment cores (Griffin et al., 2021; Larsson, 2003; Rademacher et al., 2021) provides an opportunity to automate detection and subsequent measurement of annual ring widths. Previous attempts at identifying ring boundaries have had variable and sometimes impressive success (Conner et al., 1998; Fabijańska et al., 2017; Fabijańska & Danek, 2018; Martinez-Garcia et al., 2021; Resente et al., 2021) but the only deployable method is limited to three-dimensional images (Martinez-Garcia et al., 2021). Here we present a new supervised Mask R-CNN and processing pipeline available as a Docker container. The CNN was trained and validated on RGB images of cross-dated microtome-cut conifers and has a post-processed polyline precision and recall rate of, respectively, 0.948 and 0.987 compared to cross-validated human detection. Detection is enhanced by overlapping and rotating tiles during prediction.

## Materials and Methods

### Dataset preparation and description

We formed two separate datasets of increment core samples of Norway Spruce (*Picea abies*) from one location each in Slovakia and Romania. Both plots are part of an extensive network of old growth primary forests (www.remoteforests.org), but from distinct genetic populations (Bernhardsson et al., 2020). All samples were surfaced using a core microtome (Gärtner & Nievergelt, 2010). Preparation with the microtome creates a smooth surface that leaves the cells of the wood open, in contrast with sanded samples which fills the cells with sawdust (Gärtner & Schweingruber, 2013), obscuring cell boundaries but increasing the contrast in ring boundaries.

Cores were imaged using a Zeiss Axiozoom V16 with a mechanical measuring stage. A series of 1.329×1.329 mm tiled images were collected at 70x magnification with 10% overlap. Tiles were exported as TIFF images at 33% of their original resolution because larger images ran into the integer limits of the TIFF format. Tiled images were stitched in FIJI, a well supported open source image analysis software (Schindelin et al., 2012) using the Grid/Collection Stitching Plugin (Preibisch et al., 2009).

### Training and validation data

The core images from Slovakia were cut into short segments along the long axis. Images were hand-annotated using the free online tool CVAT (Sekachev et al., 2020). Ring boundaries were annotated as a polyline then transformed into polygons by adding a buffer of 30 pixels in all directions from the original line. Each polygon was loaded by the Mask R-CNN algorithm as an individual binary mask.

Initially, the model was trained on a dataset consisting of 1705 images with 4092 annotated rings, where 179 were unseen validation images with 419 rings. In order to improve model performance on problematic sections of cores that might have been underrepresented in our original training set, we tested our first model on a new set of 67 samples of core images from our processing pipeline. These were visually inspected and regions expressing detection problems were further added to both the training and the validation set and the model was retrained. Thus, the final training dataset consisted of 2601 images and contained 8042 rings. The validation set for the final model consisted of 556 unseen images with 2596 rings as a validation dataset.

### Real-world test set

As a test of the real world application we analyzed 34 full-length increment core samples for a total of 4,355 rings, from trees growing in Romania using the CNN with the weights derived from the Slovakian samples. Prior to this analysis we cross-dated and measured the Romanian samples using Coo Recorder (Larsson, 2003; Maxwell & Larsson, 2021), dating and measurement quality was verified in COFECHA (Holmes et al., 1986). Only cross-dated samples with good image quality were used to verify CNN results.

### Model and training

An open source implementation of Mask R-CNN (Abdulla, 2017; He et al., 2017) with ResNet 101 backbone was modified for instance segmentation of tree-ring boundaries. In order to reduce training time, the model was initiated with weights pre-trained on the ImageNet dataset (Russakovsky et al., 2015). We chose an input size of 1024 by 1024 and all three RGB channels, thus all images were automatically resized to these dimensions. Image augmentation was used to increase variation in the training dataset. Namely, we used horizontal and vertical flip, rotation in the range of 90 degrees in both directions, cropped out by 30% or cropped in by 5%, further Gaussian blur, brightness, and saturation were also varied. Per image, between one to five of these augmentations were randomly applied.

As we used pre-trained weights, we started training only head branches, which were randomly initiated, for 15 epochs with a learning rate of 1e-3. Then for 45 epochs, we trained the upper 4 layers of the network and all layers for another 140 epochs. Finally, we reduced the learning rate to 1e-4 and continued training for another 200 epochs. We used weight decay of 1e-4 and learning momentum of 0.9. When the training finished we used weights from the epoch that reached the highest mean average precision (mAP) on the validation set. We used python 3.6 and TensorFlow 1.14 with Keras 2.2 on a Nvidia Quadro RTX 6000 of a computational cluster at the Vienna BioCenter.

### Pre- and postprocessing to improve model accuracy

Combining multiple detection masks can lead to improved performance of a model (Oskar Vuola et al., 2019). In order to further improve the accuracy of our model and reduce mis-detections (such as in a corner of a frame or certain orientations), we applied the following image processing steps. First, the top and bottom 17% (of image height) of the full-size core image was cropped out to remove the sloping edge of the increment core. A sliding window with 75% overlap was then used to extract squared sections, so that each section of the image is analyzed multiple times in slightly different local contexts. Tree-ring detection was then run on each of these squared sections in normal orientation, but also 45° and 90° rotations. Resulting binary masks from all sliding window positions and all rotations were added up in a two-dimensional mask of the size of the original full-size core image. Values ranged from 0 to the maximum number of overlaps of detected masks. Furthermore, ring polygons were extracted only from areas on the mask where at least three detections overlapped and only ring polygons that were bigger on either axis than 20% of image height were used. Finally, centerlines for ring polygons were calculated using an open source algorithm based on the Voronoi diagram (https://github.com/ungarj/label_centerlines), which was considered to be the final ring boundary prediction. Measurements were extracted as minimal distances between the predicted ring lines. Final measurements for each core are exported as JSON and POS files, the latter is compatible with Coo Recorder, making incorporation in familiar workflows easier.

### Model validation

To evaluate the performance of the trained model on the validation dataset we used both established metrics from deep learning as well as functional characterization against the assessment of a trained dendroecologist. This is more representative because the rings, which are in reality polylines, are evaluated as polygons using the established metrics.

We selected the best training weights from validation as the average precision over Intersection-over-Union (IoU) for thresholds from 0.5 to 0.95 with a step size of 0.05, following the COCO dataset challenge (https://cocodataset.org). The selected set of weights was evaluated in detail based on precision, recall and IoU (Table 1, Figure 1).

**FIGURE 1.**
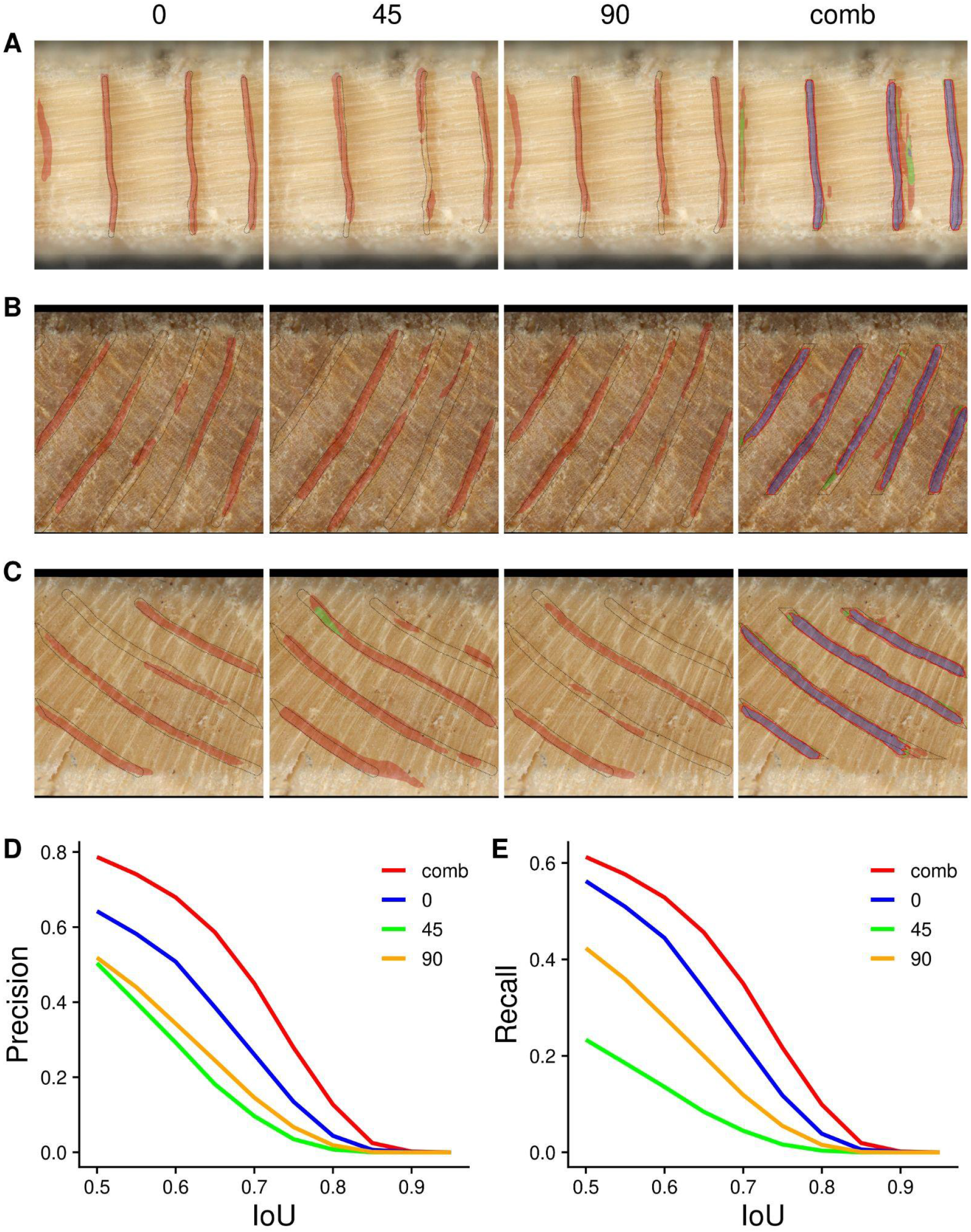
A - C) Example images showing detection at different rotations of the image (0°, 45° and 90° degrees) and combined mask as a result of processing. Red area indicate detection by one mask, green two and blue three and more. Detected mask is delineated by the red line and ground truth by the dashed line. D) Shows precision and E) recall per IoU levels for each rotation of image and combined mask as calculated on the validation dataset.

**TABLE 1.** Evaluation of the final model for ring detection on the computer validation dataset. Common metrics’ results on the validation dataset are reported for normal orientation (0°), 45° and 90° as well as the combined mask after image processing.

|  | 0 | 45 | 90 | combined |
| --- | --- | --- | --- | --- |
| **mAP** | 0.29 | 0.14 | 0.20 |  |
| **AP50** | 0.67 | 0.42 | 0.56 |  |
| **TP** | 1460 | 606 | 1099 | 1590 |
| **FP** | 813 | 597 | 1020 | 431 |
| **FN** | 1136 | 1990 | 1497 | 1006 |
| **Precision** | 0.64 | 0.50 | 0.52 | 0.79 |
| **Recall** | 0.56 | 0.23 | 0.42 | 0.61 |
| **IoU** | 0.55 | 0.42 | 0.49 | 0.64 |

Detected masks were considered a true positive if their IoU with a ground truth mask was higher than 0.5, otherwise they were counted as false positives. Ground truth instances that did not have higher IoU than 0.5 with any predicted mask were considered false negatives. These values were calculated for the whole validation set. Precision and recall were consequently extrapolated from these values. We calculated the mean IoU per image and these were averaged to generate an IoU value for the whole dataset.

### Real world testing on full core data

We inspected 34 images generated as part of the output which showed the detections overlaid on images of the sample, for a total of 4,355 rings. Each image was checked for false and missed detections. When possible these were attributed to one of several sources. These sources can be generalized as relating to 1) sample preparation, such as issues attributable to the physical preparation of the sample such as improper surface preparation, 2) sample acquisition issues relating to the imaging and processing of samples such as focus or stitching issues or 3) biological responses inherent to the sample itself such as an injury response, cracks or micro rings. In addition, we identified a number of detections which included at least part of an actual ring but were in some way flawed.

#### Partial

Detections that covered less than half of the ring without a clear mitigating factor.

#### L rings

Detections which began on a ring boundary but bent towards another shape

#### U rings

Detections which began on one ring boundary made a turn and finished on a separate ring boundary

#### Jumped rails

Detections which began on one ring boundary but ended on another ring while staying parallel to the ring boundaries

#### Post-Processing Error

Boundaries which should have been identified as rings based on the combined detections across rotations and overlapping views but were not classified as rings.

Because the above issues with detections did not fall on the true ring boundary but still indicated the presence of a true ring, we calculated precision and recall two ways. The strict values calculate precision as the true detections divided by the number of predicted detections and recall as the true detections divided by the number of true ring boundaries as calculated by the software cofecha total plus one (because it counts full rings, not boundaries). The permissive values include the flawed detections above as “true”.

### Comparison of measurements

After comparing ring detections to the actual rings we compared the measurements obtained by the minimum distance outputs of the CNN. To compare rings we used Coo Recorder to modify the POS files generated by the CNN to ensure that the ring width measurements referred to the same ring. Modification was kept to a minimum but was essential for a one to one comparison for each ring as measured manually and by the CNN and is comparable to a real world workflow.

## Results

### Training and validation set testing

Our ring detection model achieved the best performance at epoch 186 with mAP of 0.29. We decided to use these weights with detection confidence higher than 0.5. Precision, recall and IoU of the trained model reached 0.64, 0.56, and 0.55 respectively on the validation dataset without our pre- and post-processing. Applying image processing steps further improved all these metrics (precision = 0.79, recall = 0.61, IoU = 0.64; see Table 1).

### Real world testing

The real world test of the CNN showed that the model works on samples from other populations. The new (2601 trained images) CNN weights did not improve precision relative to the old (1705 trained images) for either the permissive or strict values at 0.969/0.948 (permissive/strict) for the new weights vs 0.977/0.955 for the old weights. Recall in both the permissive and strict forms was improved slightly 0.987/0.987 for the new weights compared to 0.985/0.984 for the old weights.

Most missed detections (77%) were uncategorized as they often occurred amid many similar looking rings, which are hard for the CNN to predict because of the 30 pixel buffer imposed on annotated lines in training. Those that we were able to classify were most often attributed to a rotated core (55%) (showing the sides of the tracheids instead of the tops) either because of a branch or poor sample preparation.The other clear causes of missed detections were at crack boundaries (27%) or at the edge of the image (18%).

Improper detections were more common and occured for a variety of reasons. Most often these related generally to the sample (84%), specifically, false detections in the latewood, which was not explicitly trained in the model, accounted for 24% of false detections. Another relatively common source of error was false detections either at the incomplete ring just prior to the bark or detections in the bark itself (19%). Other causes of false detections were less common but did occur, 7% of errors related to image acquisition and 2% to sample preparation.

Regression modeling demonstrates that the CNN derived measurements are strongly predictive of the human measurements with an adjusted R^2^ of 0.987. The CNN measurements were slightly smaller than the human measurements, with a model intercept of 0.049. This is consistent with the methods of deriving measurements based on minimum distance (Figure 2).

**FIGURE 2.**
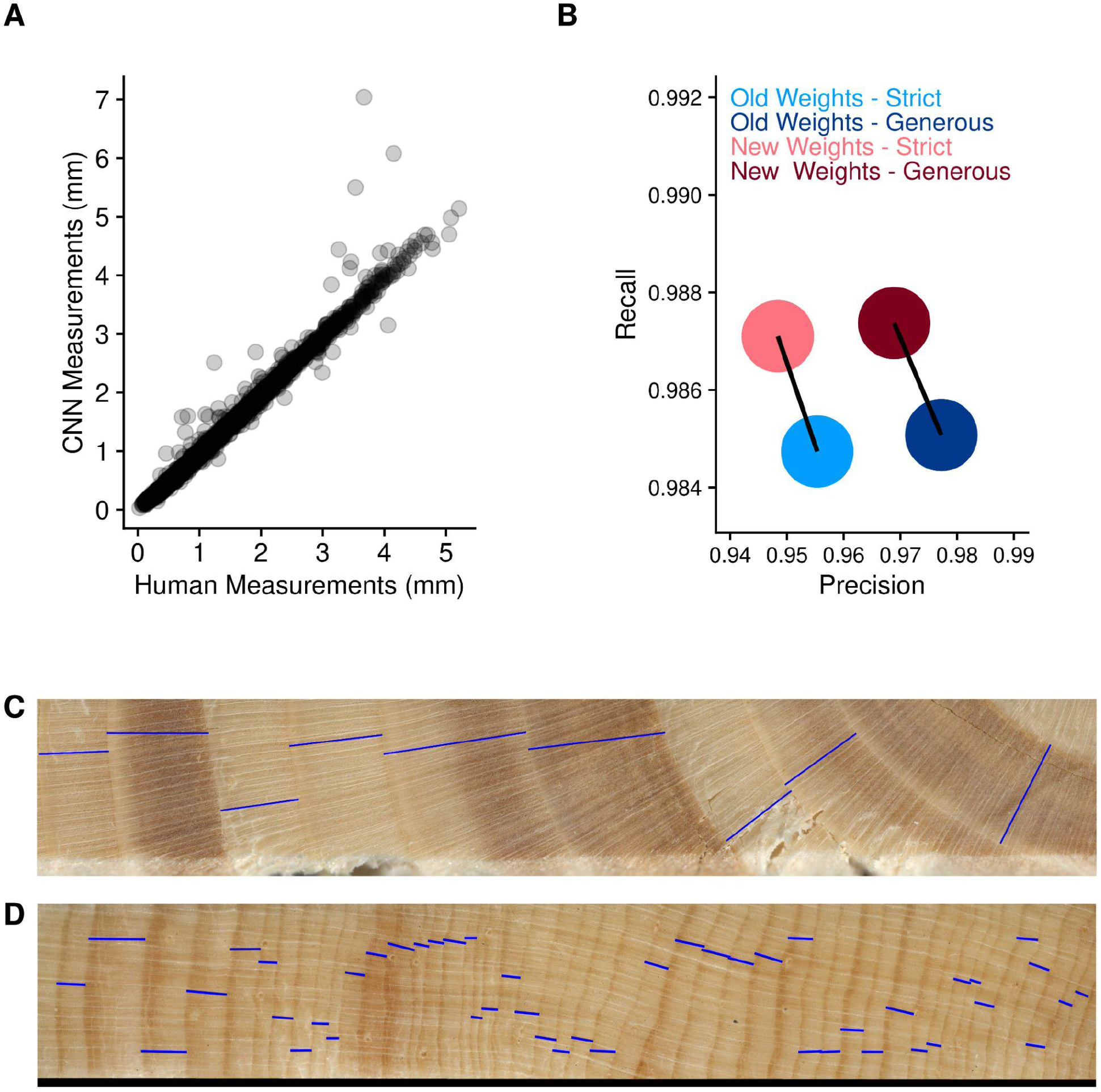
A) Comparison of tree-ring width measurements between the expert dendrochronologist (A. A.) and our neural network model on the real world samples from our core processing and imaging pipeline. B) Precision and recall evaluated by dendrochronologist on the real world sample. C-D) Examples of real world core samples. The blue lines indicate measured ring-widths as detected by our model using minimum distances between detected rings.

## Discussion

Image pre- and post-processing improves the performance of the ring detection model. Despite including image augmentation in the training, the model predictions differ between the three orientations of the same image. Overall performance is better in normal orientation and decreases with 90° and 45° rotation (see Table 1, Figure 1). However, in some cases such as when rings are angled as they approach the center of the tree, detection is better for the 45° rotation. By combining masks, we predict ring boundaries better than we can in any given orientation (see Figure 1B and C). Additionally, using overlap of at least three masks removes certain mis-detections that are detected only in certain orientations or close to the edge of the image (see Figure 1A).

Adding mis-detections back into the training set only marginally changed real-world accuracies, with recall slightly higher but decreasing precision marginally. Microrings are the largest remaining source of missed detections and this is likely due to the overlap in buffered area for these rings during training. While smaller prediction windows using higher resolution could mitigate these problems, there is a tradeoff with computational time. It is important to note that some rings that were counted as false positives on the validation images were successfully detected at the correct position of a ring and the center line would lay correctly at the ring boundary but that these rings were smaller than the threshold of IoU of 0.5 and thus were not counted as true positive. From a practical perspective, these detections would be correct and that might be reflected in higher precision on real tree core samples.

Our tree-ring CNN performed well on both the annotated validation data set and the real world test of full length cores.The CNN successfully identified rings spanning much of the variation in conifer ring appearance including relatively wide rings with reaction wood and relatively narrow rings with latewood density fluctuations (Figure 2C and D). This is a powerful tool that can greatly cut down on the processing time to generate ring-width measurements while maintaining the benefits of image based measurements (Griffin et al., 2021; Maxwell & Larsson, 2021). Though trained only on *Picea abies*, the consistent structure of conifer wood means it should be easily deployable across other conifer species (Fabijańska et al., 2017; Martinez-Garcia et al., 2021; Resente et al., 2021). We consider this CNN to be a valuable tool for efficiently generating data that can reduce analysis time considerably, especially in large datasets or for potential reanalysis of old unmeasured datasets.Though our CNN performed well with 95% of its detections correct and correctly identifying 99% of true rings, these errors still prohibit an unsupervised use of the generated data and requires an experienced dendrochronologist to oversee crossdating (Holmes et al., 1986). The CNN’s outputs include JSON files of all detections as well as position (.POS) files that are compatible with Coo Recorder (Larsson 2003). The POS files make the CNN results easily editable through Coo recorder and the recent development of new, open source image analysis programs such as WIAD (Rademacher et al. 2021) or Dendro Elevator (https://dendro.elevator.umn.edu/) provide additional ways to display both images and annotations.This software combined could provide a freeware open source image analysis pipeline.

## Acknowledgements

The data collection was possible thanks to Miroslav Svoboda, Martin Mikoláš and others behind the Remote forest database. Ethan Stewart provided useful advice about neural network implementation. Miguel Vallebueno, Laura Morales, Julia Reifler were great help in the field.

## Conflict of Interest statement

Authors declare that there is no conflict of interests.

## Data Availability

The application for tree-ring measurements is freely accessible as a Docker container at quay.io/repository/treeringgenomics/image-processing. Full code that was used for preparation of this application is also available freely and open source at GitHub (https://github.com/Gregor-Mendel-Institute/TRG-ImageProcessing/tree/TRG-ImageProcessing_Clean/CoreProcessingPipelineScripts/CNN/Mask_RCNN). Training and validation dataset and their annotation files will be available as a Dryad dataset.

## Author Contributions

K.S. conceived of the project as well as supervised all the work. M.P. selected the algorithm, performed training and developed the pre and post-processing pipeline. A.A. performed all dendrochronological analyses and quality control as well as supervised all work requiring dendrochronological expertise. P.H. designed and prepared the Docker container. K.S, M.P., A.A., L.W. and S.S. participated in data collection as well as annotation of training and validation dataset. M.P. and A.A. led the writing of the manuscript. All coauthors contributed to the preparation of the manuscript.

## Notes

### Competing Interest Statement

The authors have declared no competing interest.

## References

Abdulla, W. (2017). Mask R-CNN for object detection and instance segmentation on Keras and TensorFlow. GitHub Repository. https://github.com/matterport/Mask_RCNN

Bernhardsson, C., Wang, X., Eklöf, H., & Ingvarsson, P. K. (2020). Variant calling using NGS and sequence capture data for population and evolutionary genomic inferences in Norway Spruce (Picea abies). In bioRxiv (p. 805994). https://doi.org/10.1101/805994

Conner, W. S., Schowengerdt, R. A., Munro, M., & Hughes, M. K. (1998). Design of a computer vision based tree ring dating system. 1998 IEEE Southwest Symposium on Image Analysis and Interpretation (Cat. No.98EX165), 256–261.

Crawford, D. (2021). Earlywood Vessel Area Analysis of Quercus macrocarpa Tree Rings at the Cedar Creek Ecosystem Science Reserve in Minnesota (D. Griffin (ed.)) [University of Minnesota]. https://www.proquest.com/dissertations-theses/earlywood-vessel-area-analysis-em-quercus/docview/2572604884/se-2

Fabijańska, A., & Danek, M. (2018). DeepDendro – A tree rings detector based on a deep convolutional neural network. Computers and Electronics in Agriculture, 150, 353–363.

Fabijańska, A., Danek, M., Barniak, J., & Piórkowski, A. (2017). Towards automatic tree rings detection in images of scanned wood samples. Computers and Electronics in Agriculture, 140, 279–289.

Fricker, G. A., Ventura, J. D., Wolf, J. A., North, M. P., Davis, F. W., & Franklin, J. (2019). A Convolutional Neural Network Classifier Identifies Tree Species in Mixed-Conifer Forest from Hyperspectral Imagery. Remote Sensing, 11(19), 2326.

Fritts, H. C. (1971). Dendroclimatology and Dendroecology. Quaternary Research, 1(4), 419–449.

Gärtner, H., & Nievergelt, D. (2010). The core-microtome: A new tool for surface preparation on cores and time series analysis of varying cell parameters. Dendrochronologia, 28(2), 85–92.

Gärtner, H., & Schweingruber, F. H. (2013). Microscopic preparation techniques for plant stem analysis. Verlag Dr. Kessel.

Griffin, D., Porter, S. T., Trumper, M. L., Carlson, K. E., Crawford, D. J., Schwalen, D., & McFadden, C. H. (2021). Gigapixel Macro Photography of Tree Rings. Tree-Ring Research, 77(2), 86–94.

Gudelek, M. U., Boluk, S. A., & Ozbayoglu, A. M. (2017). A deep learning based stock trading model with 2-D CNN trend detection. 2017 IEEE Symposium Series on Computational Intelligence (SSCI), 1–8.

He, K., Gkioxari, G., Dollár, P., & Girshick, R. (2017). Mask r-cnn. Proceedings of the IEEE International Conference on Computer Vision, 2961–2969.

Holmes, R. L., Adams, R. K., & Fritts, H. C. (1986). Tree-ring chronologies of western north America: California, eastern Oregon and northern Great Basin with procedures used in the chronology development work including users manuals for computer programs COFECHA and ARSTAN. https://repository.arizona.edu/handle/10150/304672

Housset, J. M., Nadeau, S., Isabel, N., Depardieu, C., Duchesne, I., Lenz, P., & Girardin, M. P. (2018). Tree rings provide a new class of phenotypes for genetic associations that foster insights into adaptation of conifers to climate change. The New Phytologist, 218(2), 630–645.

Kido, S., Hirano, Y., & Hashimoto, N. (2018). Detection and classification of lung abnormalities by use of convolutional neural network (CNN) and regions with CNN features (R-CNN). In 2018 International Workshop on Advanced Image Technology (IWAIT). https://doi.org/10.1109/iwait.2018.8369798

Larsson, L. A. (2003). CooRecorder: image co-ordinate recording program. Cybis Elektronik & Data AB, Saltsjöbaden, Sweden.

Levanič, T (2007). Atrics - A New System for Image Acquisition in Dendrochronology. Tree-Ring Research, 63(2), 117–122.

Martinez-Garcia, J., Stelzner, I., Stelzner, J., Gwerder, D., & Schuetz, P. (2021). Automated 3D tree-ring detection and measurement from X-ray computed tomography. Dendrochronologia, 69, 125877.

Maxwell, R. S., & Larsson, L.-A. (2021). Measuring tree-ring widths using the CooRecorder software application. Dendrochronologia, 67, 125841.

Oskar Vuola, A., Ullah Akram, S., & Kannala, J. (2019). Mask-RCNN and U-net Ensembled for Nuclei Segmentation. arXiv E-Prints, arXiv:1901.10170.

Preibisch, S., Saalfeld, S., & Tomancak, P. (2009). Globally optimal stitching of tiled 3D microscopic image acquisitions. Bioinformatics, 25(11), 1463–1465.

Primicia, I., Camarero, J. J., Janda, P., Čada, V., Morrissey, R. C., Trotsiuk, V., Bače, R., Teodosiu, M., & Svoboda, M. (2015). Age, competition, disturbance and elevation effects on tree and stand growth response of primary Picea abies forest to climate. Forest Ecology and Management, 354, 77–86.

Rademacher, T., Seyednasrollah, B., Basler, D., Cheng, J., Mandra, T., Miller, E., Lin, Z., Orwig, D. A., Pederson, N., Pfister, H., Wei, D., Yao, L., & Richardson, A. D. (2021). The Wood Image Analysis and Dataset (WIAD): Open-access visual analysis tools to advance the ecological data revolution. In Methods in Ecology and Evolution. https://doi.org/10.1111/2041-210x.13717

Resente, G., Gillert, A., Trouillier, M., Anadon-Rosell, A., Peters, R. L., von Arx, G., von Lukas, U., & Wilmking, M. (2021). Mask, Train, Repeat! Artificial Intelligence for Quantitative Wood Anatomy. Frontiers in Plant Science, 12, 767400.

Russakovsky, O., Deng, J., Su, H., Krause, J., Satheesh, S., Ma, S., Huang, Z., Karpathy, A., Khosla, A., Bernstein, M., Berg, A. C., & Fei-Fei, L. (2015). ImageNet Large Scale Visual Recognition Challenge. International Journal of Computer Vision, 115(3), 211–252.

Rydval, M., Larsson, L.-Å., McGlynn, L., Gunnarson, B. E., Loader, N. J., Young, G. H. F., & Wilson, R. (2014). Blue intensity for dendroclimatology: Should we have the blues? Experiments from Scotland. Dendrochronologia, 32(3), 191–204.

Schindelin, J., Arganda-Carreras, I., Frise, E., Kaynig, V., Longair, M., Pietzsch, T., Preibisch, S., Rueden, C., Saalfeld, S., Schmid, B., Tinevez, J.-Y., White, D. J., Hartenstein, V., Eliceiri, K., Tomancak, P., & Cardona, A. (2012). Fiji: an open-source platform for biological-image analysis. Nature Methods, 9(7), 676–682.

Sekachev, B., Manovich, N., Zhiltsov, M., Zhavoronkov, A., Kalinin, D., Hoff, B., TOsmanov, Kruchinin, D., Zankevich, A., DmitriySidnev, Markelov, M., Johannes, Chenuet, M., a-andre, telenachos, Melnikov, A., Kim, J., Ilouz, L., Glazov, N.,… Truong. (2020). opencv/cvat (Version v1.1.0) [Computer software]. Zenodo. https://doi.org/10.5281/zenodo.4009388

von Arx, G., & Carrer, M. (2014). ROXAS - A new tool to build centuries-long tracheid-lumen chronologies in conifers. Dendrochronologia, 32(3), 290–293.

Washburn, J. D., Mejia-Guerra, M. K., Ramstein, G., Kremling, K. A., Valluru, R., Buckler, E. S., & Wang, H. (2019). Evolutionarily informed deep learning methods for predicting relative transcript abundance from DNA sequence. Proceedings of the National Academy of Sciences of the United States of America, 116(12), 5542–5549.

Yang, H., & Yin, L. (2017). CNN based 3D facial expression recognition using masking and landmark features. 2017 Seventh International Conference on Affective Computing and Intelligent Interaction (ACII), 556–560.

